# A common problem in the purification of calicivirus raised by molecular sieving: balance between purification and loss of viral particles

**DOI:** 10.1101/2020.02.04.933390

**Authors:** Zehui Qu, Hongtao Kang, Chenxi Cui, Kaiwen Meng, Xinzheng Zhang, Liandong Qu, Geng Meng

## Abstract

Caliciviridae is a group of RNA viruses could lead to gastrointestinal disease in humans and acute oral or upper respiratory tract disease in felines, no efficient vaccine so far. The preparation of immunogen is a challenge to obtain a safe and efficient vaccine candidate. Molecular sieving, due to the advantage of simple and fast purifying, is considered as a very powerful tool in calicivirus purification; however, also introduce mechanical damage to the viron. In this study, we took Feline Calicivirus (FCV), a common natural pathogen in cats as well as a classic model for caliciviruses studies, as a research object. Here, with the help of the cryo-electron microscope (cryo-EM), we collected particle images in both conditions of with and without molecular sieving post-processing, the impact of molecular sieving on FCV particles was evaluated by building a flexibility evaluation system after 3D classification. The results indicated that the molecular sieving will impact the stability of P domains through increasing flexibility. In addition, we successfully built up a scoring system to describe this flexibility of FCV P-domains, which can be extended to assess the purification effect to the entire calicivirus.

## Introduction

Caliciviridae are nonenveloped, positive-stranded RNA viruses including Norovirus, Sapovirus, Lagovirus, Vesivirus and so on[1]. Norovirus and Sapovirus could lead to gastrointestinal disease in humans and not easy to study in laboratory[2]. Feline Calicivirus (FCV), due to the advantages of the safety as well as the ease to culture and replicate compared with other kinds of viruses in Caliciviridae, shows important research value and becomes a very good model for relative studies. For example, Conley etc. reported calicivirus VP2 formed a portal-like assembly following receptor engagement, firstly deeply explored the receptor engagement of the Caliciviridae as well as all of the nonenveloped RNA viruses which is from the perspective of FCV[3], shared very important information on virus entry and infection mechanism in Caliciviridae. FCV is a common natural pathogen of cats[4], some specific strains will contribute to limping disease[5], acute oral and upper respiratory tract disease or even virulent systemic disease (VSD) with high mortality rates[6, 7], doing huge harm to cats. For this reason, vaccination based on inactive or attenuated viruses plays a more and more important role in FCV prevention and control. Currently, single vaccine strains (F9 or 255) has been used for several decades, which has shown a limit to a certain spectrum of field strains[8, 9]. Despite a combination of two vaccine strains (G1 and 431) has been more effective, resent vaccines have been shown certain degree of defects in the protection effect against all strains of FCV[8-12], and developing a more effective vaccine is still needed.

Whether for virological basic research or new vaccine development, the viral purification step is very important, and inappropriate purification methods will result in changes in the virus’s own properties. It is because virus state is a key factor and will directly influence the vaccine immunization as well as storage quality, which is majorly determined by the virus purification. Various methods have been applied for vaccine purification, including ultracentrifugation and molecular sieving[13]. In addition, ultracentrifugation is widely used for virus particles purification because of its perfect separation capacity, however, because that some kind of medium is needed to implement the density gradient during centrifugation, such as cesium chloride, sucrose gradients[14], which sometimes may bring damage to the vaccine composition[15]. To solve this problem, some procedures need to be carried out to eliminate the redundant gradients. Molecular sieving has the advantage of simple and fast purifying and can be used to prepare larger amounts of protein, is often used for virus directly purification as well as the post-process procedure after ultracentrifugation to remove density gradient medium. However, mechanical damage due to gel tunnels of the molecular sieving will do harm to many viruses, decrease their infectivity.

In our experiment, to check how the mechanical damage of molecular sieving will influence the FCV virus particles, after virus purification with ultracentrifugation, we post-processed the purified samples using and not using molecular sieving respectively. Under the help of the cryo-electron microscope (cryo-EM), we collected particle images in both conditions of with and without molecular sieving post-processing. By building a flexibility evaluation system by the Fourier Shell Correlation (FSC) resolution assessment after 3D classification, we find that the molecular sieving will influence FCV particles P domains flexibility. Our work shows that the molecular sieving will impact the stability of P domains through increasing flexibility, making sense to better FCV purification as well as vaccine preparation, which also provides a new way to fast, easily and quantizably analyze the calicivirus stability.

## Materials and Methods

### Virus culture and purification

Feline calicivirus strain 2280 was propagated in Crandell Reese Feline Kidney (CRFK) cells. And the virus supernatant was collected when cytopathic effect (CPE) reaches 70 – 80%. Firstly, the cell debris was pelleted from the culture medium by centrifugation (3, 000 *r.p.m.*, 20 min at 4 °C). The virus particles within the supernatant were pelleted by centrifugation at 100, 000g for 2h at 4 °C using a Surespin630 rotor. The pellets were resuspended and dissolved in 1ml phosphate buffered saline (PBS) over night at 4 °C. Then, the virus was purified by centrifugation through a sucrose gradient (40% - 60%) using a SW-41 Ti rotor at 38,000 r.p.m. for 2 h at 12 °C. The resultant pellet was obtained between 40% and 60% layers. The de-sugared experiment was carried out with three methods respectively, one part of the pellet was sampled to molecular sieve, the second part is applied to dialysis while the left part was directly concentrated in the concentrating tube and liquid changed for 3 times. Final concentrated samples were stored at 4 °C until use.

### Negative-stain electron microscopy

FCV particles were imaged by negative-stain transmission electron microscopy (TEM). 5 μl of virus was loaded onto a freshly glow-discharged continuous carbon TEM grid for 60s. The grid was washed in two 5 μl droplets of 2% p-phthalic acide (PTA), drained and dried at room temperature. The grids were then imaged in a FEI Tecnai Spirit transmission electron microscope equipped with a EMSIS VELETA 2K*2K CCD.

### Sample preparation and cryo-EM data acquisition

After the H_2_/O_2_ glow-discharge, freshly processed 300 mesh holey carbon grids (GIG-300M-Cu-R2/2, China) were used to load purified virus sample at 85% humidity and 4°C using a Vitrobot Mark IV. After pre-blot for 10s, the grids loading virus were blotted for 3s by filter paper and flash frozen by lunging into liquid ethane at -182°C using a Leica EM GP2. Cryo-EM images were collected with 200kV Talos Arctica microscope (Thermo Fisher) equipped with GIF-K2 Summit camera (Gatan) at a magnification of 100KX in the super-resolution mode yielding a calibrated pixel size of 0.66Å, each exposure of 5.28s was dose-fractionated to 32 movie frames leading to a total dose of 30 e^-^/Å^2^. Micrograph stack files were binned and firstly subjected to gain reference. After drift correction with MotionCor2 and anisotropic distortion correction, contrast transfer function (CTF) determination and defocus estimation were performed with CTFFIND4 as implemented with Relion 3.0 used for further data processing.

### Image 2D classification and 3D classification

100 virus particles were manually picked by RELION were 2D averaged to gain reference. A total of 55856 particles were autopicked and extracted (bin 4) from 1275 micrographs in RELION. After reference-free 2D classification, 55110 particles were selected (bin 2) and subjected to 3D classification with an initial model created by starticos program. 9862 particles were selected for 3D auto-refinement to get a 2.80 Å-resolution reconstruction. However, because of flexibility, the resolution of glycoprotein shell was about 4.2 Å. The rotational, translational and CTF parameters in the last iteration of the reconstruction were extracted and adapted to JSPR format by a home-made script for unbin data. The virus particles were re-extracted without binning and the 2.80 Å-resolution map was rescaled to a pixel size of 1.32Å for further refinement by block-based reconstruction method. Two blocks (five dimers near icosahedral two-fold axis, six dimers near icosahedral-three-fold axis) were selected and refined separately to overcome the flexibility among the blocks. The final resolution of the glycoprotein shell as determined by gold standard FSC using RELION post-process is 3.6Å.

After 2D classification to exclude false-positive and damaged particles, 55,110 were picked from 1,402 images. Filtered particles were applied to rounds of 3D classification and 1,384 virions with relative distinct virus spikes conformation. Then these datasets of virions were subjected to icosahedrally averaged 3D reconstructions followed by post-processing in Relion3.0 and local resolution analysis in EMAN2. Maps were visualized using UCSF Chimera.

## Result

### Purification effect between with and without molecular sieving post-processed

After ultracentrifugation, the purified virus was sampled to denaturing SDS-PAGE, the strip of FCV coat protein (VP1) appears at around 55KD (Fig.1a), which may have been proteolyzed N-terminally[16]. We then de-sugared the FCV samples with (Fig.1b) and without molecular sieving post-processed (msFCV and nmsFCV) respectively. Firstly, loaded msFCV, nmsFCV samples to the negative-stain electron microscopy. As shown in Fig.1c and Fig.1d, molecular sieving could help greatly increase the purity compared with images without molecular sieving processed, in which, the images of nmsFCV, dark background is full of impurities with negative contrast. For excluding the accidentality of experimental operation, we also applied the dialysis to process the sample as a control, as shown in the Fig.1c, the negative contrast was decreased compared with nmsFCV particles but still apparently high compared with msFCV particles, showing that if only from the perspective of FCV purification molecular sieving is a very powerful tool.

**Figure 1.**
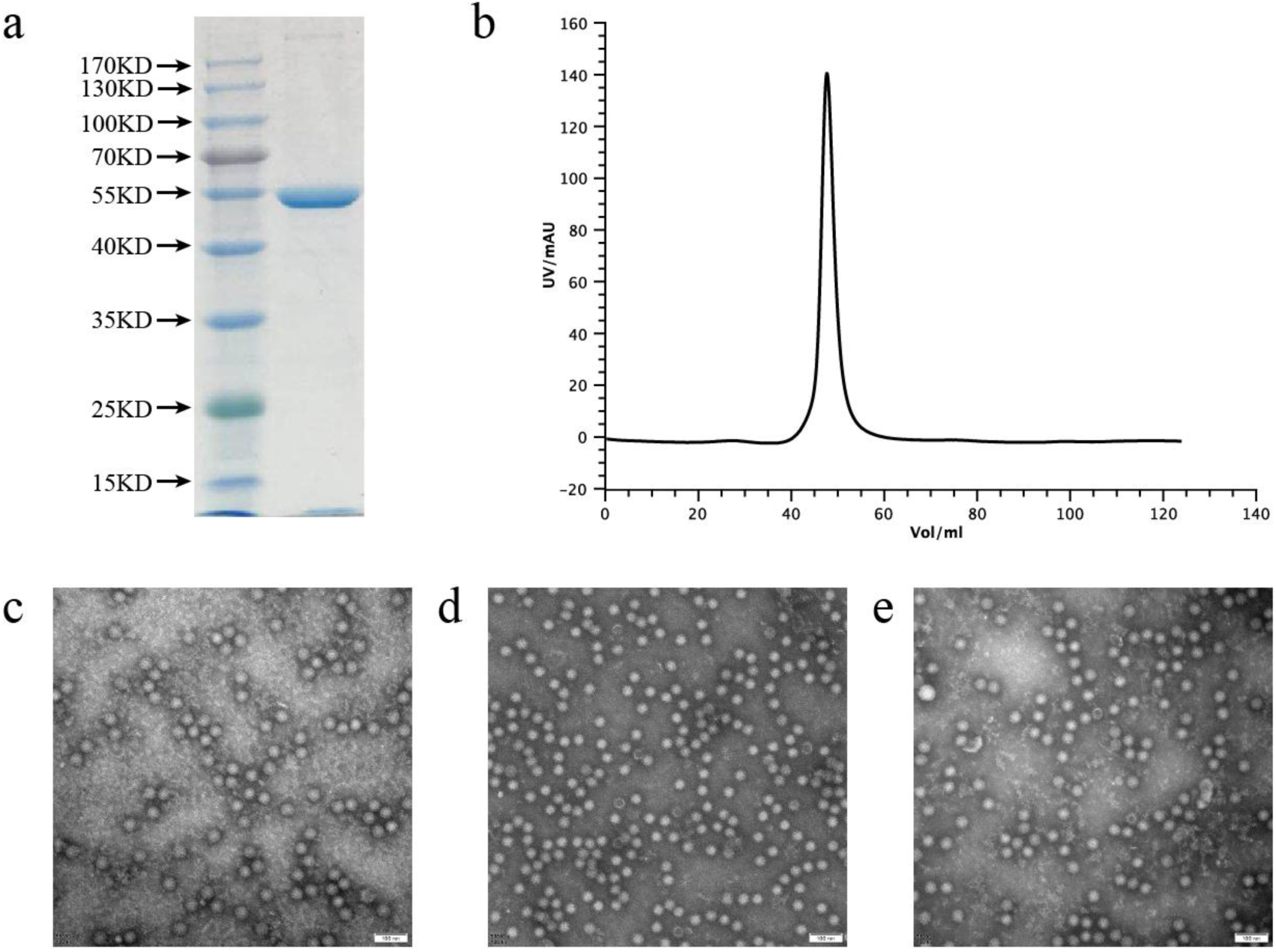
The purification effect and the negative-stain EM result. (a) The SDS-PAGE of FCV VP1 after the ultracentrifuge, the molecular weight scale is labeled on the left. (b) The molecular sieving result of the post processed VP1 after ultracentrifuge. The ordinate axis is the UV value and the abscissa axis is the volume. (c-d) The negative-stain result of the samples without molecular sieving process, with molecular sieving process and with dialysis treatment respectively. The scale was labeled at the bottom right corner (100 nm).

### Cryo-EM images comparative analysis between msFCV and nmsFCV

After the sample’s preparation, the grids loading msFCV and nmsFCV were observed under the cryo-EM, as shown in the Fig.2a. The msFCV sample shows better purity then nmsFCV sample. In the images of nmsFCV, there were more small particles, which may be VP1 that dropped off from the virion. The average defocus of msFCV images is 15061.38 angstrom, while in the nmsFCV images, the average defocus is 17934.50 angstrom. Compared with msFCV, nmsFCV images showed more discrete defocus distribution which is probably because of the more complex background noise. However, most of the the defocuses of this both samples are not badly different, showing msFCV and nmsFCV have a similar contrast and high-resolution information, which will not be influenced greatly because of the sample purity (Fig.2b). In addition, we found the msFCV was unstable compared with nmsFCV, if we do not freeze the msFCV sample in time, there will be a rapid virus disintegration (Fig.2c).

**Figure 2.**
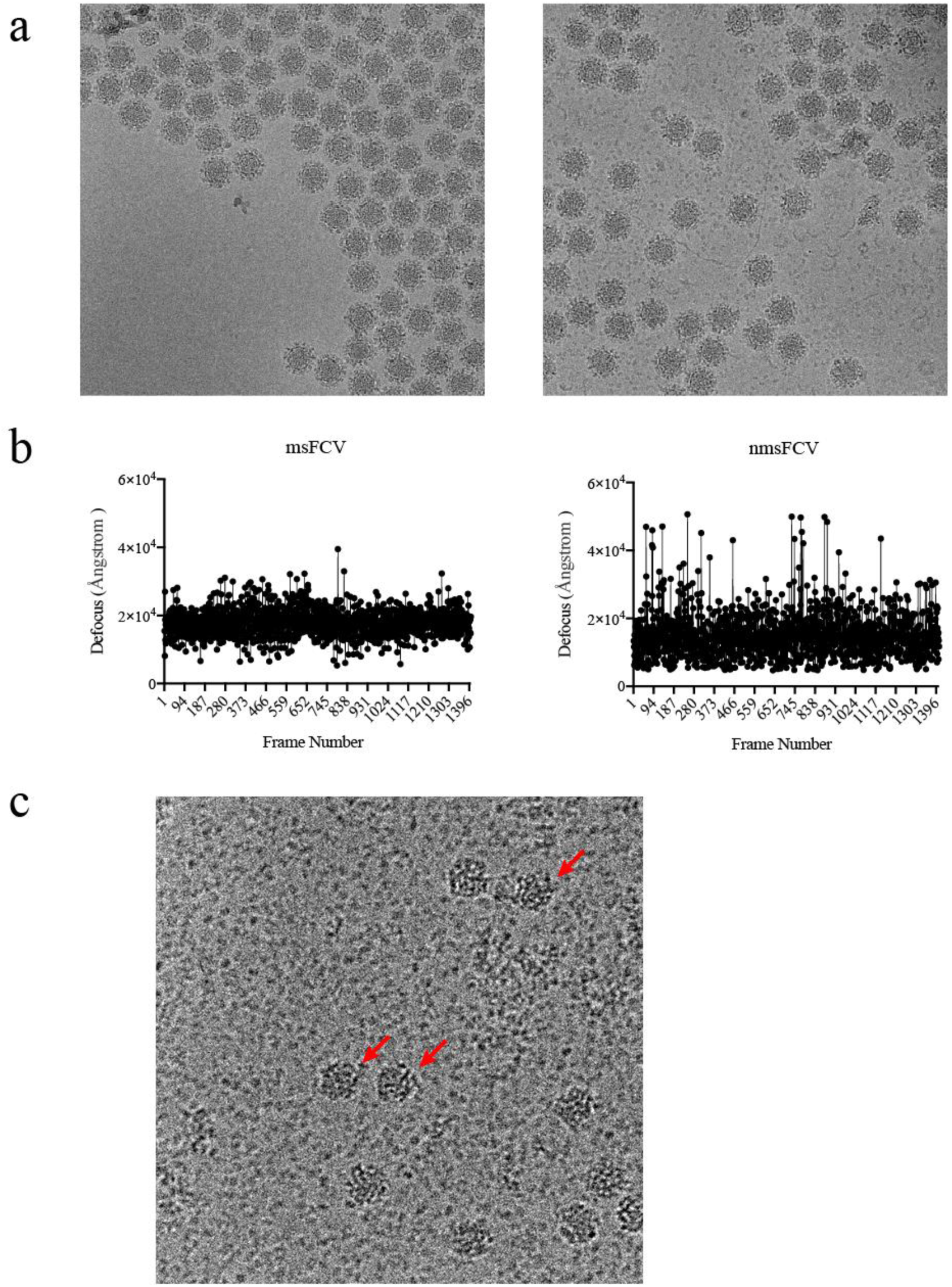
The cryo-EM analysis of the msFCV samples and the nmsFCV samples. (a) The cryo-EM images of the msFCV (left) and the nmsFCV (right). (b) The defocus distribution of the msFCV and the nmsFCV. The ordinate axis is the defocus value and the abscissa axis is the frame number. (c) The image of the degraded virus particles. The degraded particles were labeled with red arrows.

### P domains of both msFCV and nmsFCV were flexible

We firstly filtered out the bad particles through reference-free 2D classification as shown in Fig.3a. Through the noise model, we can see at the low resolution (< 13 Å), the initial sigma of the nmsFCV is slightly higher than the msFCV, and gradually becomes consistent at the higher resolution (> 13 Å), showing that the molecular sieving will help reduce the noise at the lower resolution, however does little contribution at the higher resolution compared with the samples without the molecular sieving process (Fig. 3b). After which, particles were subjected to 3D classification with an initial model created by starticos program. We firstly execute the 3D classification with default angular sampling interval (15 degrees), however we found the P-domain features of both msFCV and nmsFCV are not clear. So, we set the angular sampling interval from 7.5 to 0.1 to find the virus particles with relative stable P-domain. And interestingly, we found that there were obviously more virus particles with stable P-domain in nmsFCV particles (1384 of 55110 particles, 2.51%) compared with msFCV (171 of 21493 particles, 0.80%) (Fig. 3c).

**Figure 3.**
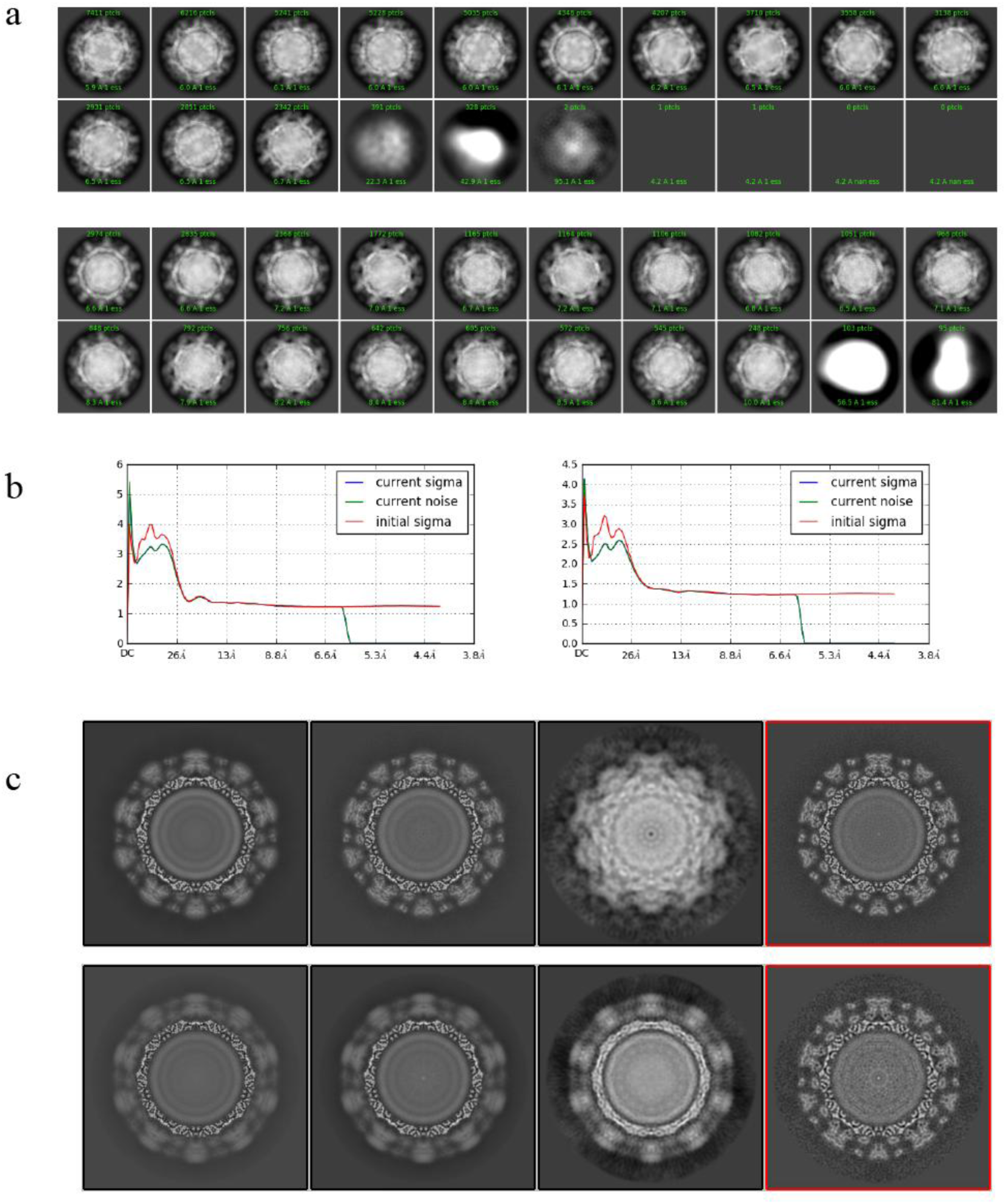
The result of the 2D classification and the 3D classification. (a) The 2D classification result of the nmsFCV (up) and the msFCV (down). Particles and the estimated resolution are labeled green in the picture. (b) The noise model of the 2D classification of the nmsFCV (up) and the msFCV (down), and the current sigma, current noise and the initial sigma are given respectively. With the abscissa axis (Resolution) and the vertical axis (sigma value). (c) The 3D classification result of the nmsFCV (up) and the msFCV (down), the P-domain stable class was selected in red box.

### Molecular sieving will do damage to FCV through increasing the flexibility of P-domains

To evaluate the effect of this molecular sieve on the P-domain flexibility of viral particles, we randomly pick out the same number (20000 particles) of msFCV and nmsFCV. Given the initial model, we applied refinement in the cryosparc[17], in which we found in the nmsFCV, there was a clearer feature of the P-domain compared with msFCV as shown in Fig.4a. It is the same with the 2D classification result, the sigma value of the msFCV is lower at the low resolution, however becomes consistent with nmcFCV at the higher resolution as shown in the Fig.4b. We have the golden standard Fourier Shell Correlation (FSC) resolution analysis of the whole particle of msFCV and nmsFCV respectively, as shown in Fig.4c. Then we have the resolution heatmap of the nmsFCV and msFCV respectively (Fig.5). We can see the S-domains of the both nmsFCV and msFCV stay the same while the P-domain of the nmsFCV is apparently higher than the msFCV showing the P domains of msFCV is much more flexible than nmsFCV, which indicating molecular sieving will do mechanical damage to the virus particle through influencing the P-domains and further increasing their flexibility. We also find the voxel number ratio (VNR) at the higher resolution (2.9 and 3.9 angstrom) will give a quantitative judgment of the damage which is brought in by the purification. In this experiment, the VNR of the nmsFCV is 0.583 while in the msFCV, the VNR is 0.330, indicating the P-domain stability negatively change with or without molecular sieving process.

**Figure 4.**
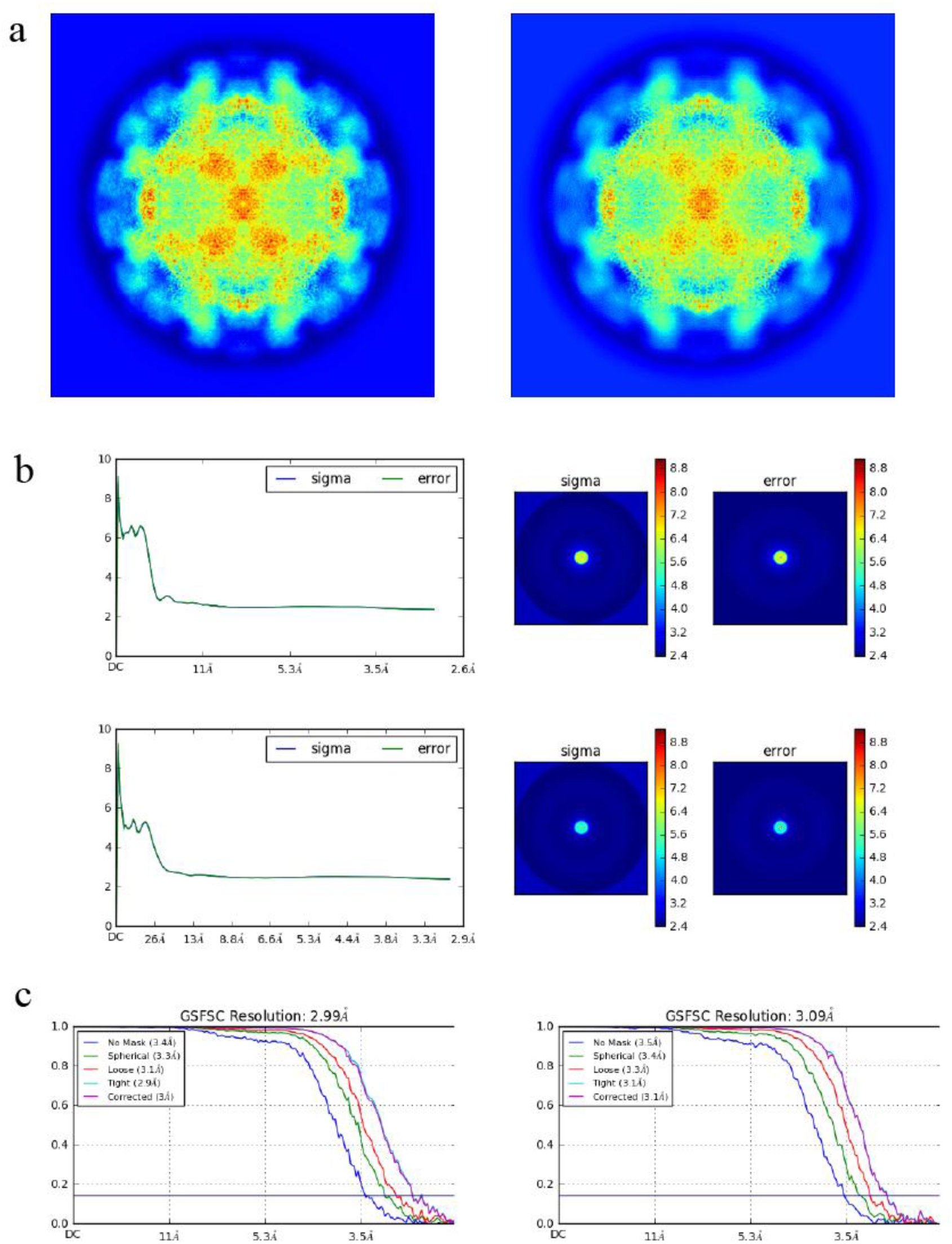
The 3D refinement result. (a) The slice picture of the nmsFCV (left) and the msFCV (right) (b) The noise model of the 3D refinement of the nmsFCV (up) and the msFCV (down). (c) The golden standard FSC curve of the nmsFCV (left) and the ms FCV (right).

**Figure 5.**
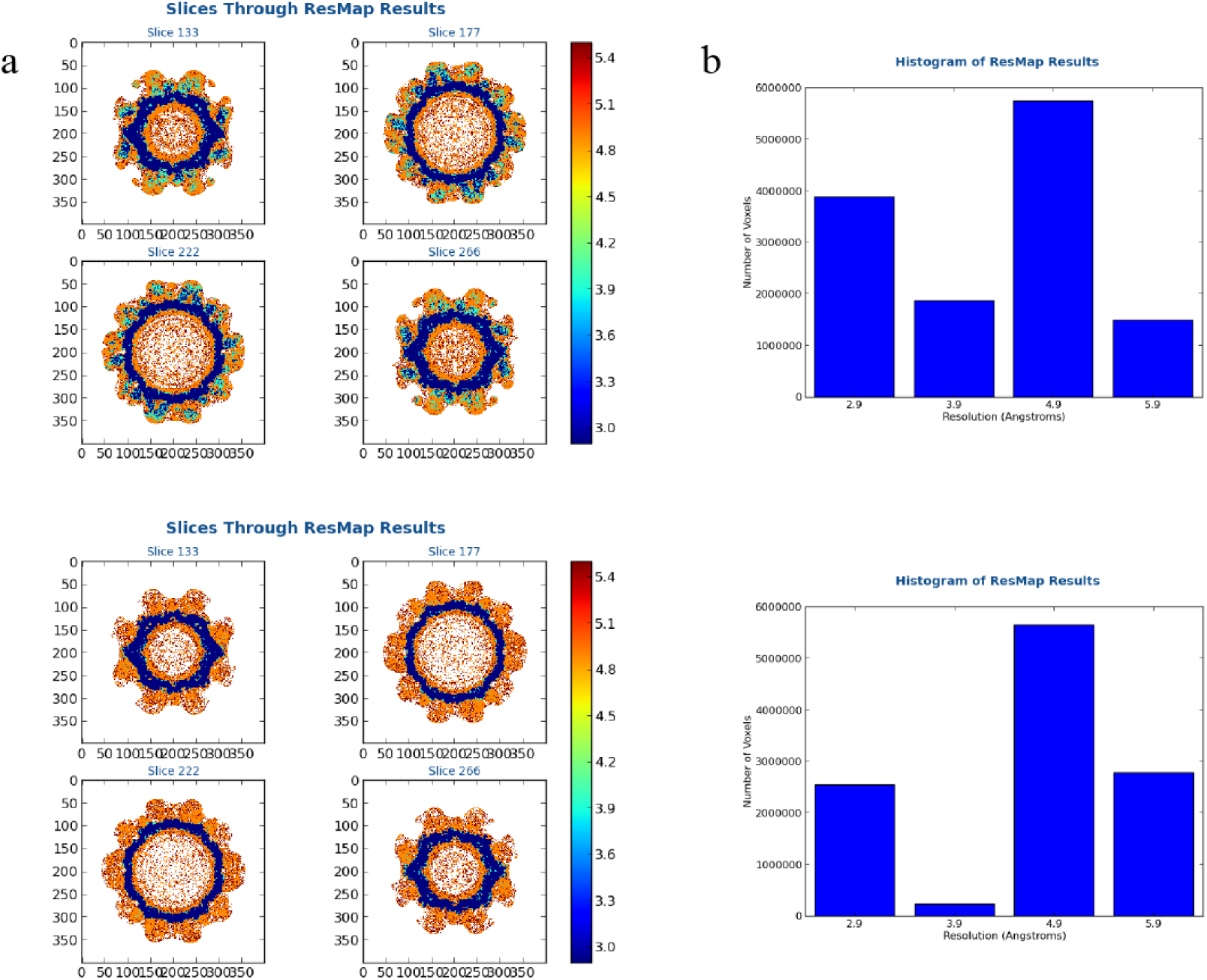
The resolution distribution after the refinement. (a) Resolution heatmap of the nmsFCV (up) and the msFCV(down), resolution scale was labeled on the right. Resolution distribution was shown as 4 gradients and colors (2.9, 3.9, 4.9 and 5.9). (b) The histogram of the voxel numbers distributed in the 2.9, 3.9, 4.9 and 5.9 gradients.

## Discussion

FCV is easy to culture and have become a very good research model to study other viruses with more virulent or more difficult to study in Caliciviridae, such as Norovirus and Sapovirus. The studies carried out around the FCV, will help us better understand and further study the other calicivirus in many of aspects, such as the vaccine development. FCV is a very harmful virus to cats, in recent years, researchers have carried out a series of work on the research of the FCV vaccine development, including inactivated virus vaccines[12], live attenuated vaccines[10], or virus like particles (VLP)[16], etc. The preparation of immunogen is a challenge to obtain a safe and efficient vaccine candidate. Virus particles of the low virulence or inactivated virus, as the most immunogenic vaccine candidate, are widely used in FCV prevention. Considering the process of vaccine preparation, after the virus grows in the cell to a sufficient titer, we collected them from the culture medium directly, which will take in a large amount of substances and impurities such as nucleic acids, proteins and lipids of the host are present in the unpurified immunogen solution. If these impurities were not processed in time, will become the heat source causing adverse reactions to the vaccinated individuals[18]. For this reason, immunogenic purity becomes the prerequisites to satisfy the vaccine safety and effectiveness. Molecular sieving or size exclusion chromatography (SEC) is a chromatographic method which separated the molecules by the size, widely applied in larger molecules purification, very suitable for virus or VLP[19]. However, molecular sieving can bring mechanical injury to some kinds of viruses. FCV is a virus with P-domains forming the ‘spi e li e’ conformation [20], which may be influenced by the gel channel. The electron microscopy analysis will help us reveal how the molecular sieving will impact on the FCV through the intuitive image information, and provide a quantified system to evaluate the balance between purification and loss of viral particles.

In our study, we find the molecular sieving is with not only excellent purification capability but also very good purification efficiency for FCV virus particles compared with other two methods mentioned, however, after molecular sieving processed the msFCV particles will become unstable and even disintegration when not prepared into cryo-samples in time. After the 3D-classification, we discovered that in the both nmsFCV and msFCV images, most of the particles were with very flexible P-domains, and only small part of particles were stable, which may be influenced not only by the gel tunnel of the molecular sieving but also the ultracentrifugation or the concentration process. However, there were significantly larger number of stable particles in nmsFCV images. We built up a scoring system to describe this flexibility of FCV P-domains, indicating that although nmsFCV’s images had a dirtier background, nmsFCV particles S-domains (core zone) could reach a similar resolution with msFCV. nmsFCV P-domains’ score is much higher than msFCV, showing that the molecular sieving will destroy the stability of the FCV particles through influencing the P-domains flexibility. This study also tells us that the flexibility of the P-domains is very important for the stability of the calicivirus, reveals a common problem in the purification of calicivirus. In addition, we believe that the flexibility scoring system established in this experiment can be extended to assess the purification effect to the entire calicivirus, lays the foundation for how to better improve the vaccine titer of cailivirus or better maintain the stability of calicivirus vaccine.

